# Divergent Behavioral Responses to Resource Limitations by Honey Bees, Bumble Bees, and Mason Bees

**DOI:** 10.64898/2026.07.20.739566

**Authors:** Lindsie M. McCabe, Kelsey K. Graham, Byron G. Love, Jonathan Berenguer Uhuad Koch, Craig Huntzinger, Diana L. Cox-Foster

## Abstract

Resource limitation is a central ecological phenomenon shaping pollinator behavior, reproduction, and community dynamics. Many studies have looked at these interactions, but few have forced competition and explored species-specific responses. Here, we experimentally evaluated behavioral and reproductive responses of honey bees (*Apis mellifera*), bumble bees (*Bombus impatiens*), and mason bees (*Osmia bruneri*) in controlled single and multispecies foraging environments. Across all treatments, each species exhibited distinct forms of behavioral compensation when exposed to increased interspecific competition. *Apis mellifera* reduced the number of foraging events in mixed species cages yet showed variation in colony growth. *B. impatiens* increased foraging effort when co-foraging with other species, but this heightened activity did not translate into increased colony growth. *O. bruneri* maintained consistent foraging effort across treatments but exhibited reduced reproduction and a marked shift in floral host use, switching from preferred species (*Phacelia tanacetifolia* and *Melilotus alba*) to less-preferred alternatives (*Collinsia grandiflora* and *Trifolium incarnatum*) when competing with social bees. Regardless of compensatory behaviors, all three species demonstrated reduced reproductive success under competitive conditions. Our findings underscore the importance of evaluating interspecific competition when placing managed bees in natural or semi-natural habitats to avoid inadvertently stressing wild or managed bee populations and taking into consideration species-specific responses in addition to community level responses to competition.

## Introduction

Resource availability is a fundamental ecological phenomenon shaping the behavior, distribution, and fitness of organisms across taxa (Tilman 1982, Holt and Bonsall 2017, Filipiak and Weiner 2017, Jeavons et al. 2022). The availability of food, nesting sites, or water are scarce and/or unpredictable, and consumers often exhibit behavioral shifts that enhance their ability to locate, acquire, and defend these limited resources (Auer et al., 2020, Abrahms et al., 2021, Duchenne et al., 2020). Such behavioral plasticity can influence community structure, alter interspecific interactions, and drive evolutionary trajectories. Among pollinators, access to floral resources determines individual survival, colony growth, and reproductive output, making them especially sensitive to changes in resource availability (Roulston and Goodell 2010, Vaudo et al. 2015). When resources become scarce, whether due to natural variability (i.e. weather), habitat degradation, or elevated forager densities, pollinators often exhibit compensatory behavioral responses aimed at maintaining energetic balance (Nagano et al. 2023, Treanore et al. 2025). Such shifts can precede detectable declines in abundance or fitness, underscoring the importance of identifying the behavioral mechanisms underpinning competitive interactions.

Competition in pollinators for limited resources generally occurs through two main pathways: interference competition and exploitative competition. Interference competition involves direct interactions such as aggression or displacement at flowers. However, in many pollinator systems, interference is relatively rare (Brittain et al. 2013, Graham et al. 2018, Rasmussen et al. 2021), whereas exploitative competition, indirect depletion of shared floral resources, has been found to affect foraging intensity, floral choice, and reproductive performance (Thomson 2004; Goulson 2010). Additional stressors, including reduced pollen quality, asynchronous flowering, and anthropogenic pressures, may further exacerbate resource limitation and its consequences (Rankin et al. 2020, Stuligross and Williams 2020, Trunschke et al. 2024, McCabe et al. 2026). Among pollinators, the interactions between managed honey bees with wild bees and how these relationships shape positive or negative fitness between species has received major research attention (Hudewenz and Klein 2015, Paini and Roberts 2005). Despite many studies looking at this competitive interaction, no generalizations have been made, and it seems like these interactions are based on location, carrying capacity of the system, stocking rates on honey bees, and which species/traits are being measured (Mallinger et al. 2017, Pike and Rittschof 2025).

Despite many studies investigating the effects of honey bee competition on wild bees or other non-*Apis* bees, almost all studies lump bees into one of two dichotomies, honey bees or non-honey bees (Hudewenz and Klein 2013, MacInnis et al. 2023) or rather just honey bees and bumble bees (Elbgami et al. 2014, Wojcik et al. 2018,). This broad grouping overlooks the importance of species specific behavioral responses to competition. More directly, experimental additions of honey bee colonies have been shown to reduce fitness in certain native species, for instance, cavity nesting bees experienced skewed sex ratios, higher offspring mortality, and reduced nest provisioning (Prendergast et al. 2025). Individual species interactions have also shown to alter honey bee foraging and increase single visit effectiveness when non *Apis* bees co forage (Brittain et al. 2013). By lumping diverse species into two or three broad groups, honey bees, other social bees, and solitary bees, meaningful variation may be concealed in how individual species respond to competition. Recognizing species specific responses is essential for understanding and maintaining resilient, functionally robust bee communities.

In this experiment, we explicitly forced interspecific competition in semi field cages to test species specific behavioral compensation and fitness outcomes for honey bees (*Apis mellifera* L), bumble bees (*Bombus impatiens* Cesson, 1863), and mason bees (*Osmia bruneri* Cockerell, 1897). We tested species-specific response as evidenced through their behavior and fitness (via reproduction or colony growth) with and without competition. To do this we examined bee-bee interactions, changes in floral selection, foraging patterns, and reproductive output in floral restricted cage environments. By establishing mixed-species pollinator communities in semi-controlled environments, we measured changes in nesting success, pollen use, and interspecific interactions. We predict that *A. mellifera* and *B. impatiens* would increase foraging effort under resource depletion but would be buffered from negative outcomes. Social bees, specifically *A. mellifera* and *B. impatiens*, can rely on labor reallocation and social homeostasis to buffer short-term stress (Seeley 1997; Donaldson-Matasci & Dornhaus 2012; Perry et al. 2015). In contrast, solitary bees, specifically *O. bruneri*, lack colony-level buffering and must instead adjust at the individual level, often through changes in floral preference or reductions in nest provisioning (Hudewenz & Klein 2015; Iwasaki et al. 2018; Eckhardt et al. 2014). Our second prediction, therefore, is that *O. bruneri* would exhibit decreased reproduction due to their solitary nature (i.e. no worker caste) which subjects them to the inability to adjust foraging efforts. However, in response to a decrease in preferred floral resources we expect to see *O. bruneri* switch hosts to less preferred plants.

## Methods

### Study System and Floral Resources

This study was conducted at the USDA ARS Pollinating Insect Research Unit (PIRU) experimental garden located on the Utah State University campus in Logan, Utah, USA over three years: 2020, 2021, and 2022. Trials took place inside portable screened cages (6.1 x 6.1 x 1.8 m) over cultivated row crops of flowering plants, using three species of bees: honey bees (*Apis mellifera*), bumble bees (*Bombus impatiens*), and mason bees (*Osmia bruneri*). Year one (2020) did not yield analyzable data because we found that unlike *O. bruneri*, which we have successfully reared in cages provisioned with a single floral species (Lacy phacelia), this method was insufficient to sustain honey bees and bumblebees (Williams and Christian 1991, Sprague et al. 2016). The following two years (2021, 2022) nine plant species from six families (Table 1) were selected to span a wide range of taxonomic diversity, flowering phenologies, and floral characteristics to provide a diverse supply of pollen and nectar throughout the growing season for all three bee species. All plants were sown from commercially available seed in rows such that each cage had an even distribution of flowers (totaling three rows of each plant species per cage), received irrigated water throughout the trials and were weeded as necessary. A total of fifteen cages were set over the rows in three replicates. Each replicate consisted of five cages randomly assigned to each bee species alone (three cages) or with all three bee species combined (two cages). Trials began when it was determined that an adequate supply of bloom (∼ 30% bloom) was available and ended after bloom was complete for a total of four weeks in 2021 (June 12 – July 10) and six weeks in 2022 (June 20 – August 2). Floral resource availability was not quantified in 2021, however in 2022 we collected weekly data on floral abundance and blooming periods of each plant species. To do this we estimated plant cover (as a percentage of the three rows planted per species) and bloom (percentage of plant cover in bloom), with three people recording values to calculate an average in each cage (Supplemental Figure 1).

**Table 1:** Average number of interactions in an observation day (8:00 – 16:00) per cage between and among species for single or multispecies treatments.

|  | Single | Mixed |  |
| --- | --- | --- | --- |
|  | Intraspecific | Intraspecific | Interspecific |
| *A. mellifera* | 0.2 | 0.4 | 1.4 |
| *B. impatiens* | 0 | 0 | 0.8 |
| *O. bruneri* | 1.8 | 3 | 0.6 |

### Bees

Micro honey bee colonies were started new each year using commercially available bees purchased from commercial supplier in California. Packages contained 1.4 lb (0.64 kg) of workers and a new queen were installed in Eco Bee Boxes (Saskatoon, Saskatchewan, Canada). Each micro colony consisted of three boxes (each containing six mini frames with a natural was starter strip) between a top cover and bottom board (with landing board). Colonies were allowed to establish for two weeks on a sucrose solution and a small patty of Ultra Bee artificial pollen (www.mannlakeltd.com), while also freely foraging for local resources outside cages. Prior to trials, each colony was inspected and equalized such that each contained a live queen, ten frames covered with worker bees, six frames of brood and three frames of honey/pollen. Fresh water was provided in each cage. Colonies were weighed weekly and inspected for varroa mites.

*B. impatiens* colonies were purchased from Koppert Biological Systems, Inc. (https://www.koppertus.com) and reduced in size so that there was one queen,10 workers, and their developing brood. Colony containers were installed in a larger plastic container with vent holes and an inspection port allowing the use of a borescope for periodic inspection as described in Koch et al. (2021). Colonies were weighed and inspected weekly.

*O. bruneri* were sourced locally from a population managed by PIRU for over ten years. Artificial nest blocks (12.7 x 15.2 x 15.2 cm) were constructed of foam laminates containing sixteen – 4 mm straws. Nest blocks were placed in cages inside an open container that provided protection from rain (approximately 1.5 m off the ground). A total of twenty bees (ten each male and female) were released in each cage. Completed nest straws were removed daily and replaced with empty straws to not limit nesting rates.

Behavioral responses evaluated in 2022 included interactions, visitations, and foraging rates.

### Interactions

To determine interference competition, we monitored behavioral interactions of five individuals of the study species once a week. In each cage, we observed the foraging behavior of five individual bees for each species. Foraging behavior data included which flowering species were visited, the duration spent on each flower or inflorescence, and if they touched or where in proximity to another bee (of the same or different species), as well as any post-interaction response (i.e. flew away or no response). Interactions were scored as a presence/absence, and included any individual being bumped, pushed, or moved by another individual, and the context of that interaction.

### Visitation

Visitation rates and floral preferences were calculated weekly by counting bees visits at each of the nine plant species for up to 30 minutes per sample at five times throughout the day at 08:00, 10:00, 12:00, 14:00 and 16:00. Observers would count the number of bees on each inflorescence, identifying both the bee species and the plant species, regardless of whether they were observed collecting pollen, nectar, or another behavior (e.g., grooming). To calculate floral preference, we used the selection ratio method described in Simanonok et al (2021). The average selection ratio ([number of bee visits to a specific plant species/total bee visits to all plant species within that site]/the percent cover available of that plant species) ± SE was calculated for each plant species in each cage. Plant species are considered selected for when the selection ratio is closer to one. For each bee species, overall differences between mixed and single cage treatments were tested using PERMANOVA on Euclidean distances computed from cage-level score vectors; divergence between treatment means was summarized using Euclidean, Cosine (1 − similarity), and Bray–Curtis distances. For each plant species, score distributions were compared between treatments using the Wilcoxon rank sum test with Benjamini–Hochberg FDR adjustment, and Cliff’s delta quantified nonparametric effect sizes and direction. All statistical analyses were conducted using R 4.1.0 using packages tidyverse (Wickham et al 2019), vegan (Oksanen et al 2026), effectsize (Ben-Shachar et al 2020), and philentropy (Drost 2018).

### Foraging rates and numbers

Foraging rates were monitored to detect differences in foraging time between single treatments and interaction treatments. We conducted foraging observations weekly over five time periods throughout the day, similar to visitation rates, but on different days. For each observation period an observer would track five individuals per bee species in each cage for up to five minutes maximum. The observer would select a bee that was coming out of the nest/hive and start recording how long that individual spent at each flower or inflorescence, and the total time it took them to return to nest/hive. Monitoring stopped when the bee returned to the hive/nest, stopped visiting flowers (e.g., bounced off cage walls or stayed in one location), or was lost to the observer (i.e., flew rapidly away).

To test the number of foragers we counted bee visits at each of the nine plant species for up to 30 minutes per sample at five times throughout the day at 08:00, 10:00, 12:00, 14:00 and 16:00. Observers would count the number of bees on each inflorescence, identifying both the bee species and the plant species, regardless of whether they were observed collecting pollen, nectar, or another behavior (e.g., grooming). we ran a generalized linear mixed model (GLMM) to test for differences between bee species and treatments (single vs mixed interactions). For this model we compared the number of plants visited in the 5-minute observation period by treatment and species, with cage # being a random effect. Additionally, we wanted to test if foraging changed throughout the day or if it changed throughout the season. To do this we ran six GLMMs, one model for each species for time of day and over the season. These models were all structured the same way by filtering just the data of one the three species (*A. mellifera, B. impatiens*, or *O. bruneri*) and averaging the number of visits across the season into their respective time intervals with visits as the response variable and treatment and time of day and the predictor variables, once again cage was a random effect. Likewise, for seasonally changes, visits were averaged to weekly observations and tested where visits were the response variable and treatment, and week were the predictor variables. All statistical analyses were done using R 4.1.0 using packages lme4 (Bates et al 2015).

### Reproductive output or colony growth

We measured reproductive output of all three bee species in species-specific ways. Reproduction and colony growth metrics were measured in 2021 and 2022. Weight (g) measurement was taken weekly to measure the reproductive success of the *A. mellifera* microcolonies. Weight is an indicator to colony growth we attributed increased colony growth as a “healthy” colony (Lefebvre and Pierre 2006). Additionally, we took measurements on several colony indicators, including number of live workers, number of dead brood, number of cells with honey, and the presence of a queen. At the end of the season frames were pulled out of each hive, and the number of live workers were counted, and the occupied cells were counted and categorized. We tested for differences in colony weight over time with linear regression models with weight as the response variable and treatment and year as predictor variables (H1: we predict that there will be a decrease in colony weight over time if there were limited resources available for *A. mellifera*). Since many of the colonies did so poorly, we ran zero-inflated GLM’s with Poisson distributions for the colony size metrics One of the three metrics (number of live workers, number of dead brood, and number of cells with honey) were used as the response variable in the fitted GLM model with treatment as the predictor variables. Prior to analyses we tested for zero-inflated and overdispersal to determine which models to use (H2: we predict that there will be a decrease in number of live workers, number of dead brood, and number of cells with honey if there were limited resources available for *A. mellifera*). Statistical analyses were done using R 4.1.0 using packages glmmTMB (Brooks et al 2017) and DHARMa (Hartig 2024) to test residuals.

*B. impatiens* colony reproduction was estimated by measuring weight of colony growth during the season, amount of colony organic matter in the colony at end of season, and number of open cells of male and females in the colonies. Colonies were weighed bi-weekly and queen status determined. At end of season, the colonies were culled by freezing and taken apart to determine number of nest cells, bumble bee created organic matter (bees, wax, silk), and other measures. From the deceased colonies, we measured total colony weight, weighed the total organic material produced, and the number of open cells. We fit three GLM’s with Poisson distributions for the colony size metrics. One of the three metrics: 1) colony weight (H1: we predict that there will be a decrease in colony weight over time if there were limited resources available for *B. impatiens*), 2) total organic material (H2: We predicted that total organic matter would increase with colony strength given that strong colonies tend to be larger), 3) open cells (H3: We predicted that colonies with more open cells would be healthier and large colonies, since open cells can be an indication on emerged individuals) were used as the response variable in the fitted GLM model with treatment as the predictor variables. Prior to analyses we tested for overdispersal to determine which model family to use. Statistical analyses were done using R 4.1.0 using packages glmmTMB (Brooks et al 2017) and DHARMa (Hartig 2024) to test residuals.

*O. bruneri,* nesting straws were collected at night once a week in the same data collection period as *A. mellifera* and *B. impatiens*. A radiograph image of each nest was taken to determine the number of cells laid during that week. Nest straws were returned to the same place that night, to not disturb reproduction. Furthermore, radiograph images were taken for all completed nests at the end of the study to determine the total number of cells made, and the contents of the cells. We used a Faxitron 48804N, (Faxitron Bioptics, Tuscon, AZ) to capture a radiograph of each bee nest at 28 KV for 15 seconds. To test if reproduction differed between treatments (single v mixed interactions) we fit four GLMs to examine four reproduction metrics: 1) number of cells made per cage (H1: we predict that there will be a change in the number of cells made per cage if there were limited resources available for the female *O. bruneri* to provision nests); 2) Number of cells per nest, (H2: *O. bruneri* typically provisions 3-5 cells per nest, likewise we suspect that a decrease in this number could indicate lower resource availability). 3) Number of alive adults pre-overwintering. (H3: *O. bruneri* overwinter as adult and there we predict that limited resources would limit development and therefore produce less bees that made it to adulthood); 4) Number of “pollen balls”, which is a provision mass that was fully made and either had an egg laid on it that never developed or never had an egg laid on it. (H4: We predict that an increase of pollen balls could signify a stressful environment.) One of these four reproduction metrics were used as the response variable in the fitted GLM model with treatment as the predictor variables, using a binomial distribution. Statistical analyses were done using R 4.1.0 using packages lme4 (Bates et al. 2015).

## Results

### Interactions

We found no evidence of interference competition between any of the species. A total of 1,598 floral visitations by individual bees were recorded, equally represented across species. Of those visits, only 41 interactions (i.e. bees “bumping” bees off visiting flowers) were observed (2.57% of total interactions) with 10 interactions observed in single species treatments and the other 31 interactions in the mixed species treatments (Tabel 1). 55% of the mixed species treatments interactions (17/31 interactions) were intraspecific interactions, mainly *O. bruneri - O. bruneri* interactions. While the other 48% (14/31 interactions) consisted of *A. mellifera- O. bruneri* interactions (19%) and *A. mellifera- B. impatiens* (26%) interactions where *A. mellifera* was “bumping” the other species off the flower.

### Floral Preference

Community floral preference between single and mixed changes did not significantly change for *A. mellifera* (df = 5, F = 0.55, p = 0.651) or *B. impatiens* (df = 5, F = 1.63, p = 0.414). However, the mixed and single treatments did impact floral preference for *O. bruneri* (df = 5, F = 3.51, p = 0.028). *O. bruneri* shifted from *P. tanacetifolia* (p = 0.009) to higher visitations of *T. incarnatum* (p = 0.005) and *R. sativus* (p = 0.002) in mixed cages (Figure 1), despite *T. incarnatum* being less abundant in some of the cages. Strong preference of *O. bruneri* for *M. alba* was also noted but we failed to detect a significant difference between treatments (p = 0.36). In both single and mixed cages *A. mellifera* preferred *M. alba* followed closely by *P. tanacetifolia* (Figure 1). *B. impatiens* strongest preference was *H. annuus* and then *M. alba* in both mixed and single cages (Figure 1). While the overall model was not significant, *B. impatiens* did show a small shift towards *P. tanacetifolia* (p = 0.051) when in mixed cages.

**Figure 1:**
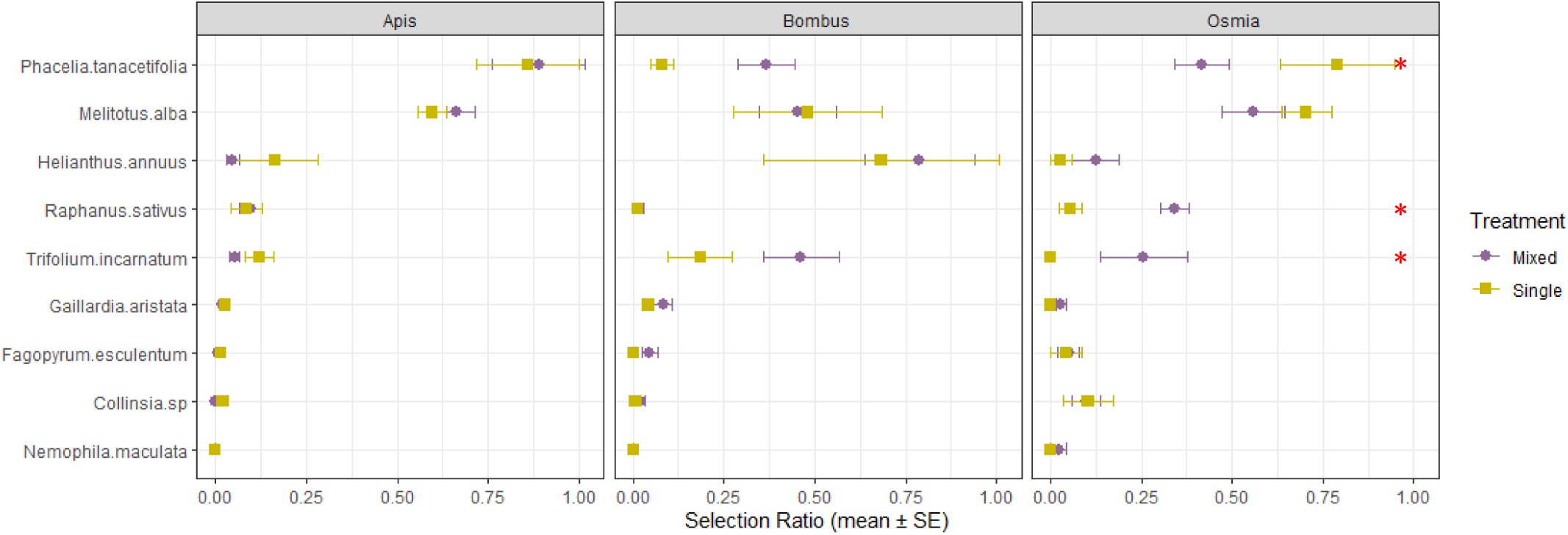
Average selection ratio (number of bee visits to a specific forb species/total bee visits to all forb species within that site)/the percent cover available of that species) ± SE for each forb species in the cage. Forb species are considered selected for when the selection ratio is closer to one. Selection ratio in the single cages is indicated by the yellow point and mixed caged are indicated by the purple point. Selection ratios where the SE does not overlap are considered significantly different from one another and indicated by a red asterisk.

**Figure 2:**
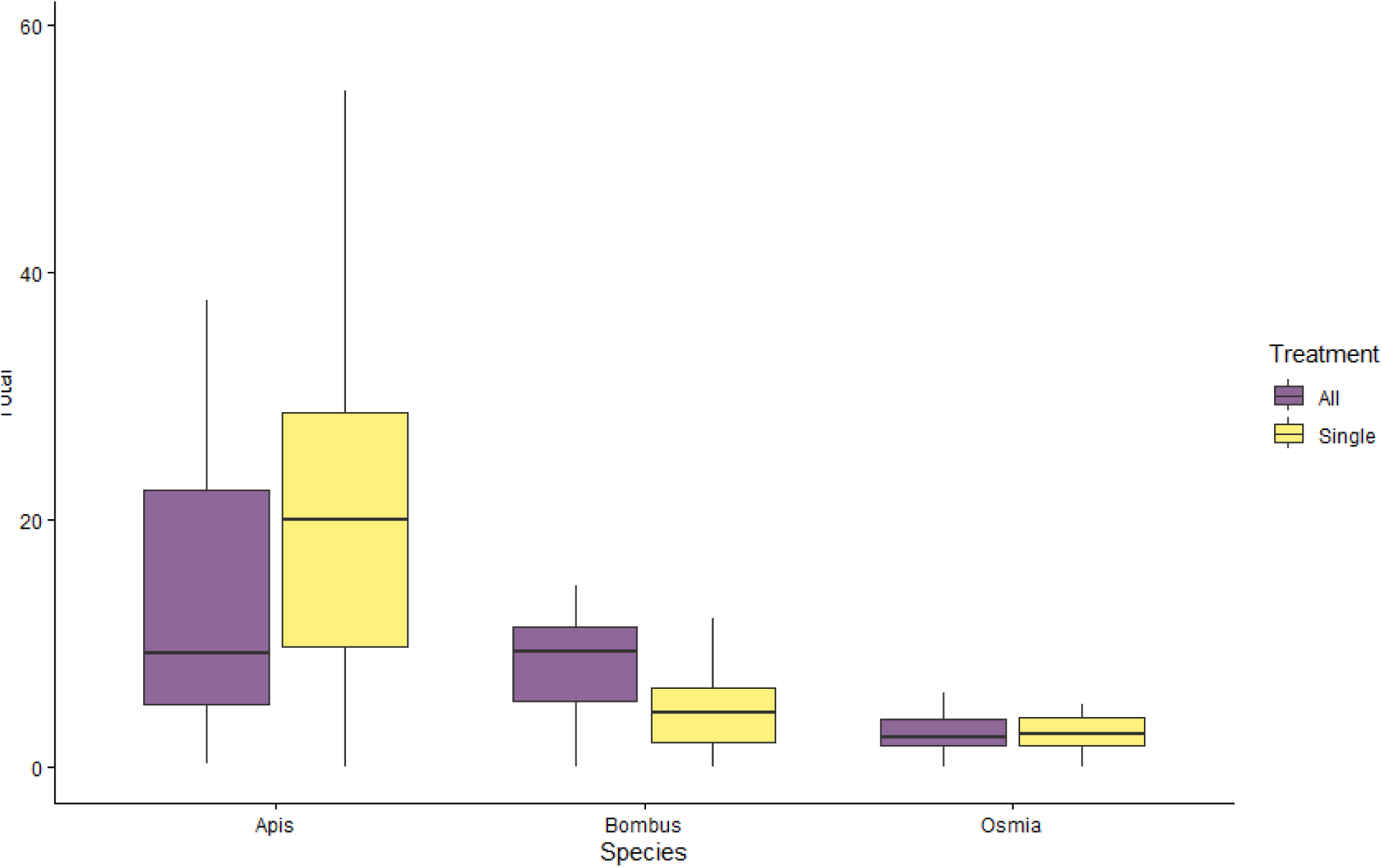
Average number of flowers visited by each species (from left to right: *A. mellifera, B. impatiens*, and *O. bruneri*) in single species cages (yellow) and mixed species cages (purple) per cage. Averaging both seasonality and time of day observations.

**Figure 3:**
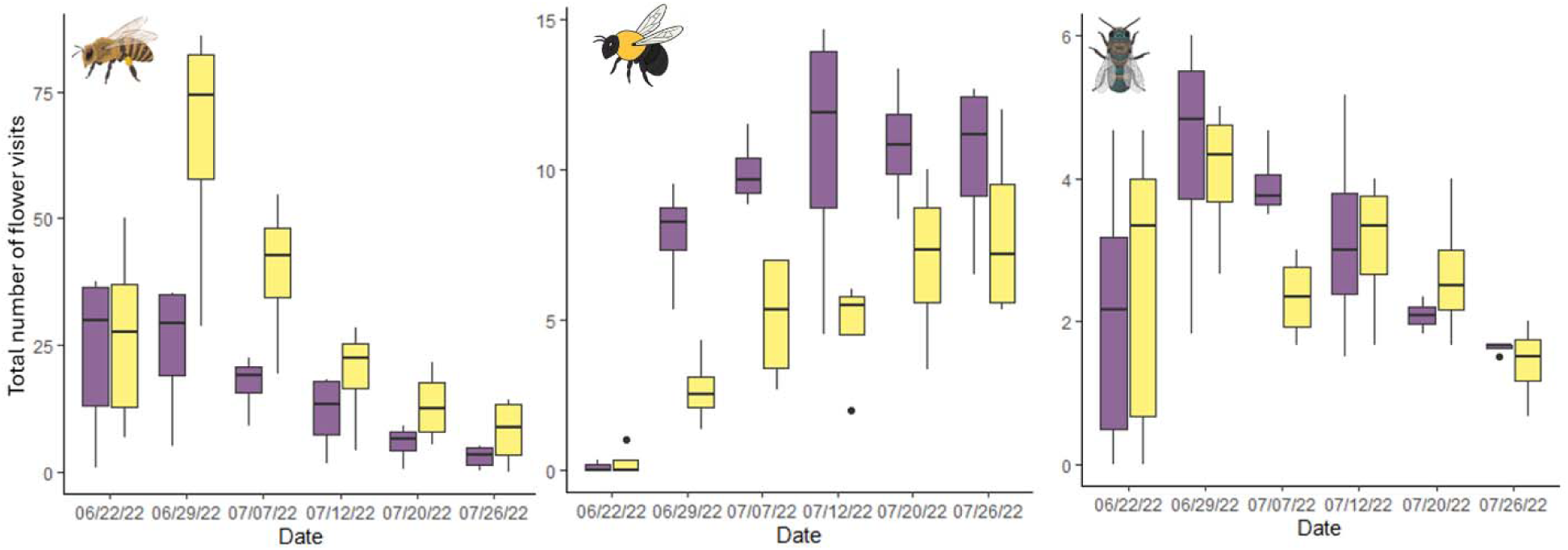
Average number of plants visited by each species (from left to right: *A. mellifera, B. impatiens*, and *O. bruneri*) in single species cages (yellow) and mixed species cages (purple) throughout the season.

**Figure 4:**
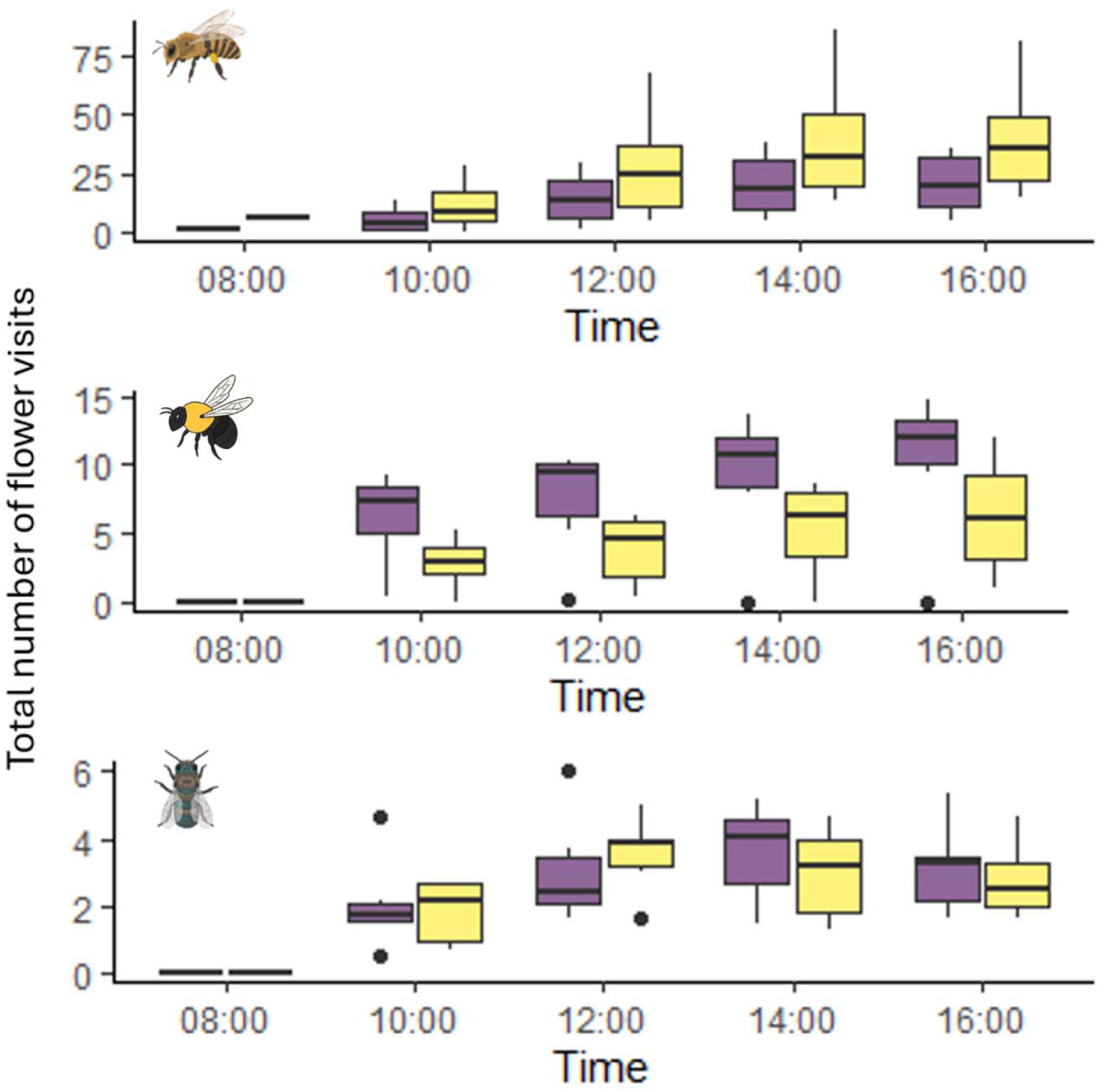
Average number of plants visited by each species (from left to right: *A. mellifera, B. impatiens*, and *O. bruneri*) in single species cages (yellow) and mixed species cages (purple) by time of day.

**Figure 5:**
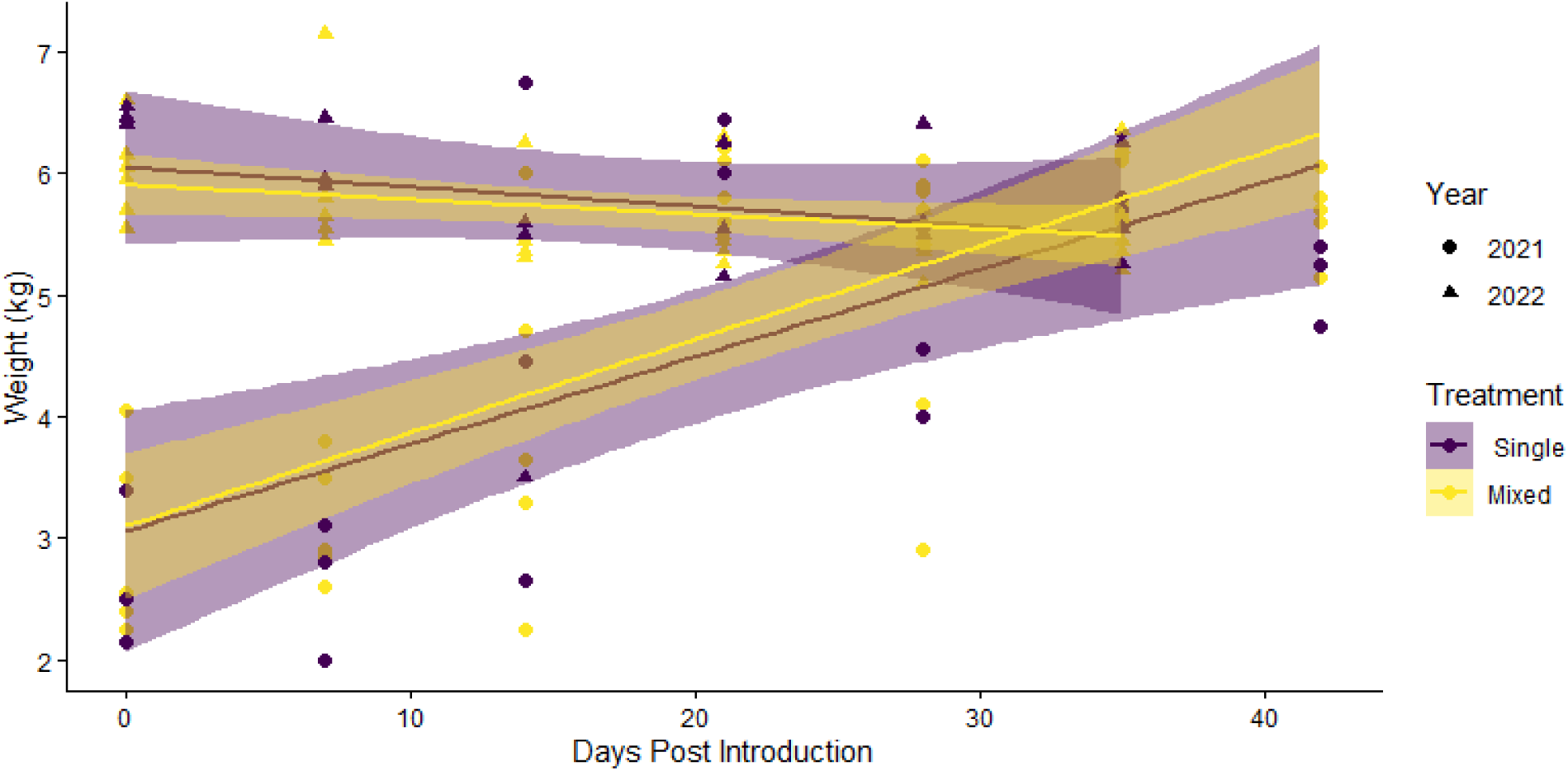
*A. mellifera* colony weight + SE over time in single species cages (yellow) and mixed species cages (purple). 2021 data is indicated by circle points and 2022 data is indicated by triangle points. Overlapping SE indicated non-significant differences between treatments or years.

### Foraging

Foraging patterns differed between species and among treatments. *A. mellifera* exhibited decreased floral visitation when placed in mixed species treatments versus when they were alone (df = 5, F = 16.72, p < 0.001). On average *A. mellifera* had 35 visitations in single species treatments and 17 visitations per cage per day in mixed species treatments. Conversely, *B. impatiens* increased their number of floral visitations (df = 5, F= 9.98, p < 0.001) with an average visitation of 4 visitations per cage per day in the single species treatments and 14 in the mixed species treatments. *O. bruneri* floral visitations did not change between treatments (df = 5, F = 0.285, p = 0.594), and on average there were 3 visitations in the single species and 3.1 visitations in mixed species treatments.

Similar patterns were observed with foraging throughout the day (∼10 hours of daytime activity observed). The number of bees foraging in a treatment cage changed based on species and time of day. The number of *A. mellifera* foragers linearly increased from 08:00 – 16:00 (df = 5, F = 7.07, p < 0.001). With the average number of foragers amounting to 10 individuals at 08:00 to nearly 40 individuals at 16:00. In cages with all three species, the number of foragers present was greater at all sampling times as compared to those cages that only contained *A. mellifera* (df = 5, F = 16.72, p < 0.001). On average, the all-species cages had 14 more *A. mellifera* foragers than in the singular species cage. *B. impatiens* foragers also increased linearly in number of foragers throughout the day; except they did not start foraging until 10:00. Average number of foragers at 10:00 was 5 and at 16:00 it was 10 individuals. Treatment cages with all species had an average of ten *B. impatiens* foragers, while bees in treatments cages with only *B. impatiens* had an average of six foragers. *O. bruneri* foraging started at 10:00 similarly to *B. impatiens.* However, rather than increasing throughout the day *O. bruneri* followed a parabolic distribution where peak foraging was between 12:00 – 14:00 with about five foragers at peak during the 14:00 hour.

### Reproductive Output

For colony weight of *A. mellifera*, the two years differed significantly in the changes in weight over the course of the season (df = 58, F= 34.3036, p < 0.001), with all colonies gaining weight in 2021 (P= 0.001), without significant differences in colony weight between treatments in singles (4.45 +/- 0.26 kg) or mixed cages (4.74 +/- 0.25 kg) (P= 0.148). In 2022, the *A. mellifera* colonies lost weight during the season (P=0.008) and competition by the three species had negative impact (P=0.023) (*A. mellifera* by self, 5.9 +/- 0.14 kg; *A. mellifera* with other species, 5.65 +/- 0.12 kg) (df=51, F=3.9035, p = 0.0032). The number of live workers, dead brood and cells with honey all showed negative impacts in the mixed species treatment compared to the single species treatment (Fig 6). Mixed species cages had 20.6 +/- 6.07 live workers at the end of the season compared to the 65.33 +/- 26.27 live workers in the single treatment (df = 5, Z= 5.395, p < 0.001). Dead brood was four times higher in mixed cages than in single species cages with 202.83 +/- 64.16 dead brood cells in mixed treatments compared to 55 +/- 11.02 dead brood cells in single species treatment (df = 5, Z = -15.73, p < 0.001). Additionally, cells with honey production also increased in single species treatments, 523 +/- 301.95 cells in single species treatment and 92 +/-45.42 cells in mixed treatments (df = 5, Z = 43.29, p < 0.001). Finally, we also noted queen presence at the end of the season. Only one queen out of the 15 colonies survived, and that was in a single species cage.

**Figure 6:**
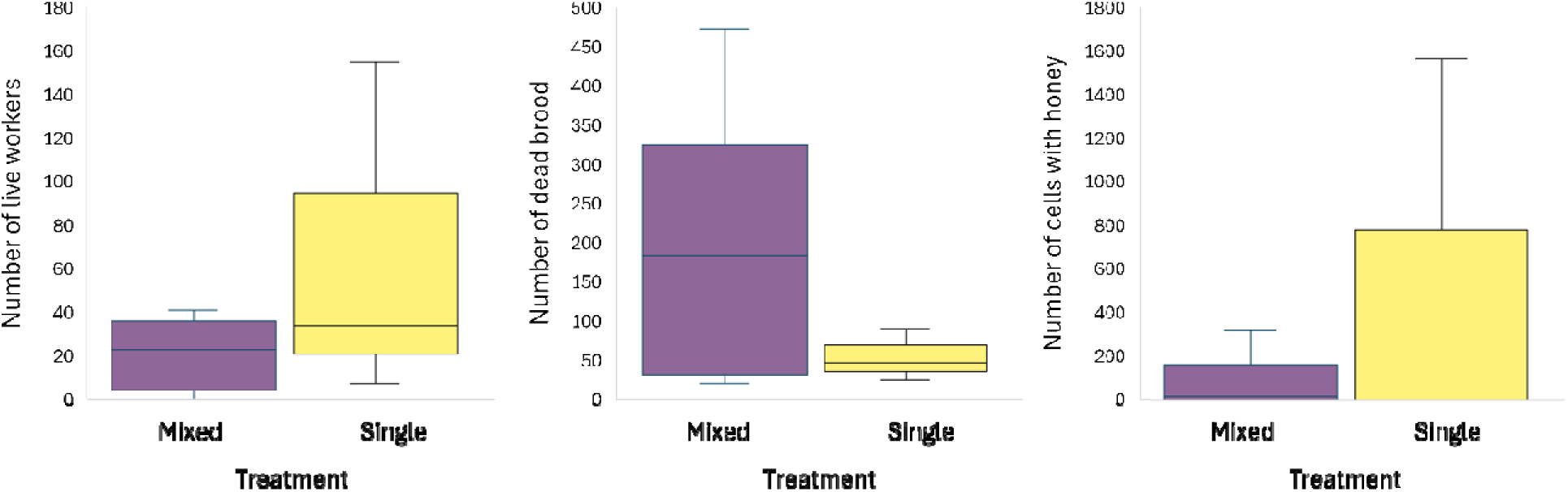
Colony success metrics for *A. mellifera* in single species cages (yellow) and mixed species cages (purple). Where colony success was measured by number of workers alive (A), number of brood produced (B), and number of cells with honey (C) at the end of the season.

**Figure 7:**
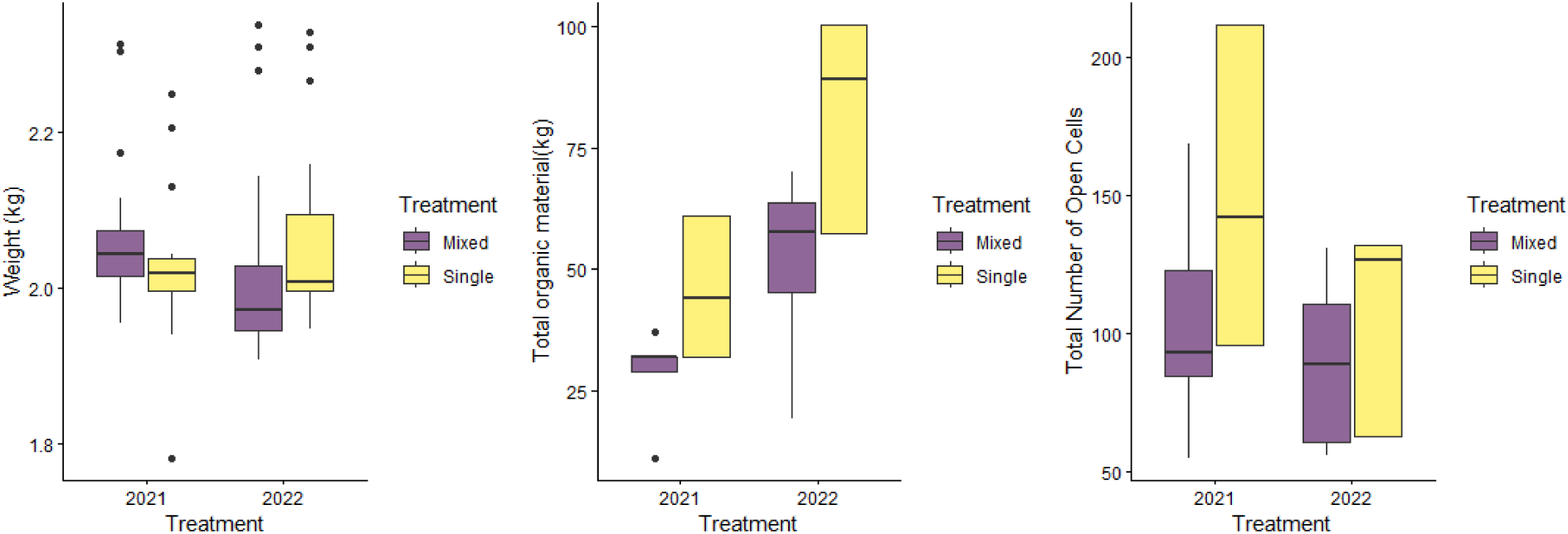
Colony success metrics for *B. impatiens* in single species cages (yellow) and mixed species cages (purple). Where reproductive success was measured by weight (A), organic material produced (B), and number of male/female cells produced (C) at the end of the season.

**Figure 8:**
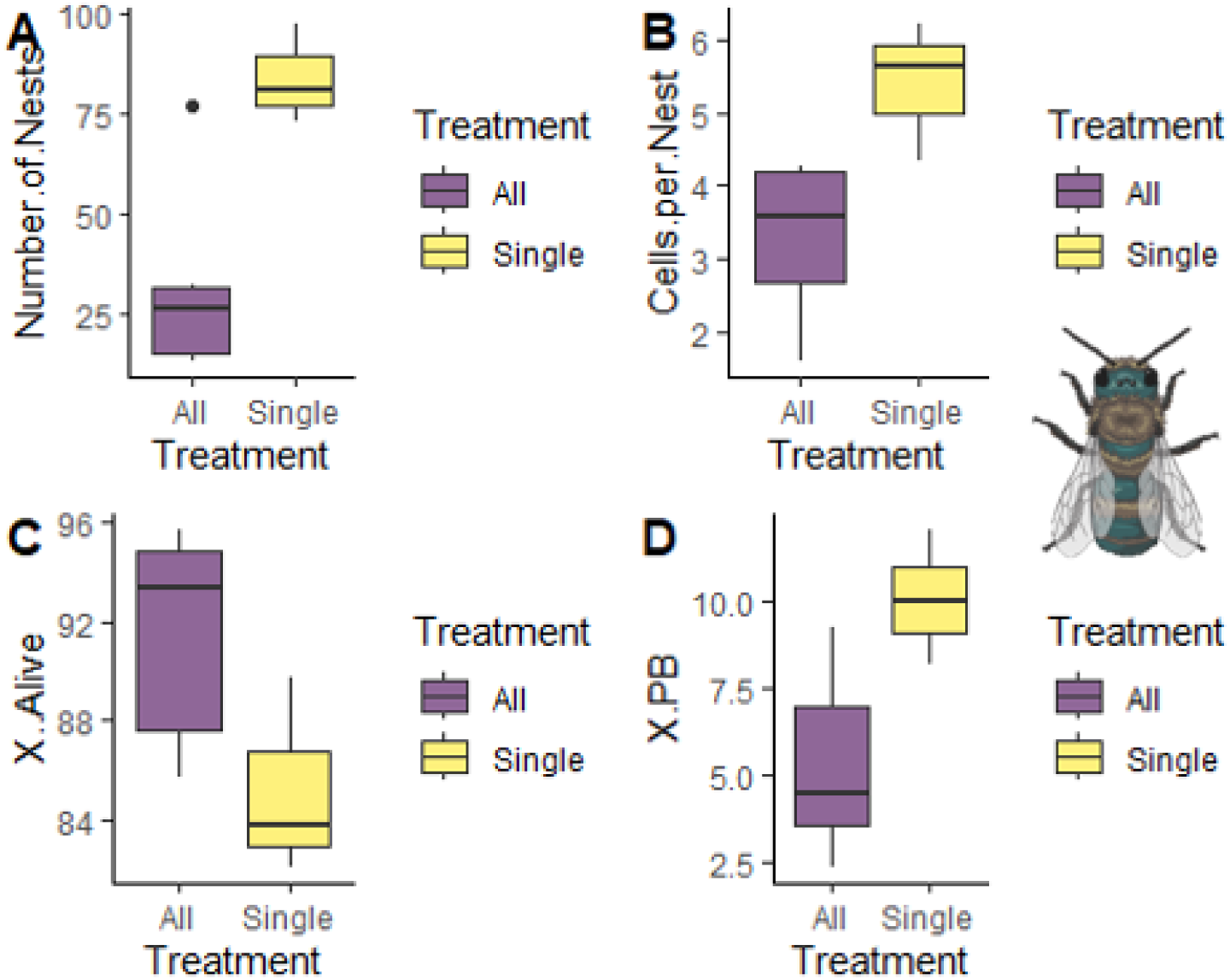
Reproductive success for *O. bruneri* in single species cages (yellow) and mixed species cages (purple). Where reproductive success was measured by A) number of nests produced per cage, B) cells made per nest and C & D) percentage of viable offspring (shown as number of eggs/provisions that made it to adulthood, and then number of pollen balls produced).

We found differences in *B. impatiens* colony weight between 2021 and 2022. We observed that colonies maintained their weights in 2021 but lost weight in 2022. Interactions with other species was only a significant effect in 2022 when looking at weight changes over time (df= 53, F=27.8197, p < 0.001), with colonies by themselves (2063.4 +/- 19.15 g) weighing significantly more than colonies with other species (2010.7 +/- 13.5 g). Measured at the end of the seasons, *B. impatiens* colonies had better growth metrics when alone than when in cages with other species. The total *B. impatiens* organic matter in each colony reflects the total productivity of the colony. In the two years, colonies by themselves had more total *B. impatiens* organic matter (62.6 g +/-3.04) than colonies with other species (37.05 g +/- 2.32) (df= 110, F= 70.07, p < 0.001), and colonies produced more organic matter in 2022 versus 2021 (p= 0.0001). Likewise, more open cells for males and females were produced in colonies by themselves (130.31 cells +/- 7.6) versus in colonies with other species (94.81 +/- 5.8) (df= 110, F= 17.125, P < 0.0001), with more open cells made in 2021 versus 2022 (p= 0.0002). In 2021, one colony by itself made three queen cells; none of the other *B. impatiens* colonies made queen cells in either 2021 or 2022.

*Osmia bruneri* experienced decreased reproduction when in cages with *A. mellifera* and *B. impatiens* (F = 12.2, p = 0.011, Figure 9). In cages where only *O. bruneri* was present, 3x more nests were made than when *O. bruneri* was in a cage with the other two species. Additionally, *O. bruneri* females made on average two more cells per nest when in cages alone than when combined with other species (F = 8.62, p = 0.026, Figure 9). Females in cages with only *O. bruneri* averaged 5.8 +\- 0.8 cells per nest while females in cages with all three species averaged 3.6 +/- 1.2 cells per nest. Both treatments experienced pollen ball and immature mortality. However, there was no significant difference in the number of pollen balls provisioned between the two treatments (F= 4.133, p = 0.081, Figure 9). However, immature mortality of *O. bruneri* was heightened in cages with all species compared to the treatment with only *O. bruneri* (F = 7.63, p = 0.028, Figure 9). On average treatments with only *O. bruneri* had 6% unexplained immature death, where all species treatments had 16% unexplained immature death.

## Discussion

Across all three focal taxa, *A. mellifera*, *B. impatiens*, and *O. bruneri*, we found evidence of species-specific behavioral compensation in response to interspecific competition, yet these adjustments were insufficient to prevent declines in reproductive output. Identifying how pollinators adjust their behavior and reproductive investment under competitive pressure is essential for predicting community stability and for designing management strategies that minimize unintended stress on wild and managed bees.

*A. mellifera* reduced their foraging activity in mixed-species cages, a pattern consistent with previous work showing that *A. mellifera* modulates foraging intensity in response to resource depletion or increased forager density (Donaldson-Matasci & Dornhaus 2012). *A. mellifera* colony growth by weight differed between years which may be due to colony health prior to release in the cages. Despite this reduction, *A. mellifera* colony growth did not differ between treatments in 2021 and showed only modest declines in 2022, which could suggest colony-level buffering (Ulgezen et al. 2024). However, the other colony size metrics showed negative impacts, where mixed species cages also had negative impacts on the colonies’ ability to build up and maintain a queen. Ultimately *A. mellifera* did better when it was just *A. mellifera* alone and not in resource-depleted environments. Additionally, it’s possible that due to poor colony growth and queen die off in both the treatments, there was not sufficient resources in the cage environments to support a honey bee colony (Banada and Paxton 1990, Abrol 2012).

*B. impatiens* exhibited the opposite pattern to *A. mellifera*. *B. impatiens* increased floral visitation in mixed-species treatments, indicating compensatory foraging to offset reduced resource availability. Similar increases in foraging effort under competitive stress have been reported in *B. impatiens* species (Thomson 2004; Goulson 2010). Yet this heightened activity did not translate into improved colony performance. Colonies in mixed-species cages produced less organic matter, fewer open brood cells, and no queen cells, suggesting that energetic constraints or reduced pollen quality limited reproductive investment. Because our colonies began the experiment with standardized worker numbers, one conclusion could be lack of gyne production in competitive environments may reflect insufficient resource accumulation to trigger the switch to reproductive output (Amslem et al. 2025). However, many of the colonies we received had already begun to produce gynes and drones, despite us requesting young colonies. This could have conflate our results with number of gyne cells produced.

*O. bruneri* showed a third compensatory strategy, *O. bruneri* maintained consistent foraging rates across treatments but shifted floral host use and reduced nesting activity. This strategy is not surprising given that *O. bruneri* is a solitary species and cannot change number of foraging females to compensate for decreased resources. Host switching from preferred species (*P. tanacetifolia*, *M. alba*) to less-preferred alternatives (*Collinsia* sp., *T. incarnatum*) mirrors patterns observed in other solitary bees under competitive pressure (Hudewenz & Klein 2015; Iwasaki et al. 2018). Because solitary bees rely on individual foraging trips to provision each brood cell, even subtle reductions in pollen quality or handling efficiency can have strong fitness consequences (Eckhardt et al. 2014). The reduced number of nests and lower cell counts per nest in mixed-species cages indicate that resource limitation directly constrained reproductive output. Additionally, the increase in unexplained immature death in the mixed species cages may indicate poor nutrition of the provision mass (Knauer et al. 2022), potentially influenced by the host switch is a less suitable/nutritious plant.

The results of this study align with broader patterns observed in other studies and highlights the subtlety of early behavioral indicators (Treanore et al 2025). None of the species shifted their foraging time of day, and interference competition was rare (<3% of interactions). Instead, changes in visitation rates, foraging intensity, and floral choice emerged as the primary behavioral signals of competitive saturation. These metrics may serve as early indicators in field settings, where resource depletion often precedes detectable declines in bee abundance or reproduction.

Some studies have reported negative effects of *A. mellifera* competition (Cane and Tepedino 2017; Page and Williams 2023a,b; Quinlan et al. 2024), while others emphasize context dependence (e.g., Paige et al 2024, Bommarco et al 2021, Pike and Rittschof 2025). In landscapes with abundant floral resources, *A. mellifera* may have minimal or no detectable impact on wild bees (Herrera 2020; Ropars et al. 2022). Some studies even document neutral or positive interactions, such as increased crop yield when *A. mellifera* and *Osmia* bees co-forage (Brittan et al. 2018; Boyle et al. 2019; McCabe et al. 2024). Our results do not contradict these findings; rather, they demonstrate that when resources are fixed and bee numbers exceed the carrying capacity, as in our cage experiment, competition becomes unavoidable, and all species incur fitness costs.

The results of this study may have important implications for placing managed bees in natural or semi-natural habitats. Even when behavioral changes appear modest, reproductive declines may already be underway and may be masked until bees exhibit measurable declines. Identifying the carrying capacity of a landscape, including temporal patterns of floral resource availability, is therefore essential for minimizing resource stressors on wild pollinators.

## Conclusions

Our experiment demonstrates that *A. mellifera*, *B. impatiens*, and *O. bruneri* bees all exhibit behavioral plasticity under competitive conditions, yet neither of the species could fully compensate for resource limitation. Behavioral shifts, reduced foraging, increased foraging, or altered floral choice, may serve as early indicators of competitive saturation, preceding measurable declines in reproduction. These findings reinforce the need to consider interspecific competition when deploying managed bees and highlight the importance of maintaining abundant, diverse floral resources to support coexistence among pollinator species.

## Acknowledgements

We greatly appreciate the dedicated work of the following, especially during the COVID years: Aletia James, Alex Fortin, Alex Foster, Alison Teague, Anne Gill, Ashley Rohde, Brock Redman, Carson Liesik, Emily Burgess, Erica Brus, Evah Peard, Harley Cragun, Jack Pugsley, Jessie Tabor, Makenna Bird, Maren Petrie, Nicole Boehme, Samuel Galt, Sierra Aston, Soliam Velez, Tien Lindsay, Timothy Olsen, and Xavier Haemmerle. Funding was provided by Project Apis M., Willard L. Eccles Foundation, USDA/ARS base funds for projects 2080-21000-019-000D and 2080-30500-001-000D.

Additionally, We thank the following for comments on our initial experimental design or for letters of support after reviewing our proposed study: Danielle Downey, George Hansen, Project Apis M.; Marla Spivak, Distinguished McKnight Professor, University of Minnesota; Margarita Lopez-Uribe, Lorenzo Langstroth Early Career Associate Professor of Entomology, Penn State University; Mace Vaughn, Scott Black, Xerces Society; Laurie Adams, Lora Morandin, Victoria Wojcik, Pollinator Partnership; Kelvin Adee, American Honey Producers Association; Joan Gunter, American Beekeeping Federation; Bret Adee, Adee Honey Farms; Jerry Hayes, Bee Culture Magazine; Dwight Wells, West Central Ohio Beekeepers Association; Randy Oliver, Golden West Bees and Scientific Beekeeping; Darren Cox, Cox Honey of Utah, LLC; Arathi Sedshari, Terry Griswold, Michael Branstetter, Theresa Pitts-Singer, Gloria deGrande-Hoffman, Kevin Hackett, USDA/ARS; Jim Cane, Vincent Tepedino, emeritus USDA/ARS.

---

**Supplemental Figure 1:**
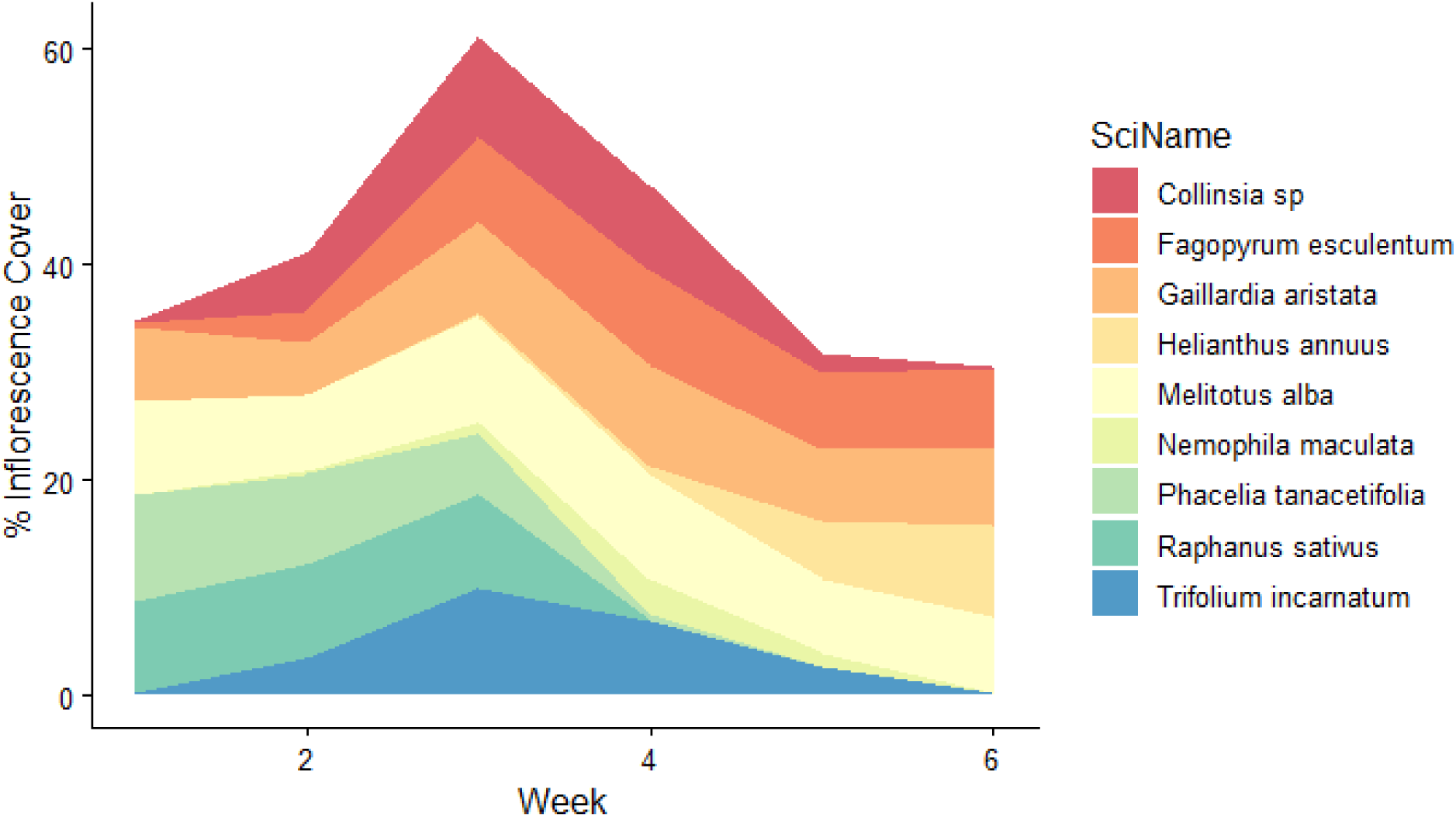
Stacked floral abundance for the duration of the study.

